# Locally coordinated synaptic plasticity shapes cell-wide plasticity of visual cortex neurons *in vivo*

**DOI:** 10.1101/249706

**Authors:** Sami El-Boustani, Jacque P K Ip, Vincent Breton-Provencher, Hiroyuki Okuno, Haruhiko Bito, Mriganka Sur

**Author notes:** Correspondence to: Mriganka Sur and Sami El-Boustani. These authors contributed equally to this work (Sami El-Boustani, Jacque P K Ip).

## Abstract

Plasticity of cortical responses involves activity-dependent changes at synapses, but the manner in which different forms of synaptic plasticity act together to create functional changes in neuronal responses remains unknown. Here we show that spike-timing induced receptive field plasticity of individual visual cortex neurons *in vivo* is anchored by increases in synaptic strength of identified spines, and is accompanied by a novel decrease in the strength of adjacent spines on a slower time scale. The locally coordinated potentiation and depression of spines involves prominent AMPA receptor redistribution via targeted expression of the immediate early gene Arc. Similar changes accompany recovery of eye-specific responses following monocular deprivation. These findings demonstrate that Hebbian strengthening of activated synapses and heterosynaptic weakening of adjacent synapses, in dendrites with heterogeneous synaptic inputs, co-operatively orchestrate cell-wide plasticity of functional neuronal responses.

**One Sentence Summary:** Arc-mediated local synaptic plasticity regulates reorganization of synaptic responses on dendritic stretches to mediate functional plasticity of neuronal responses *in vivo*.

## Main Text

Neuronal circuits in the developing and mature brain are subject to dramatic changes driven by sensory inputs (*1*,*2*) or motor learning (*3*–*5*), causing cells to modify their responses to individual inputs while maintaining a relatively stable level of activity (*6*). Global mechanisms such as cell-wide homeostatic plasticity were initially reported for stabilizing the output firing rate of a cell by uniformly scaling up or down the effective strength of all its synapses (*6*, *7*). More recent experimental evidence *in vitro* has suggested that other forms of compensatory plasticity can also act locally at the level of dendritic stretches (*8*–*13*) or even at single synapses (*14*–*16*) and that potentially confer rich functional compartmentalization within the dendritic tree (*17*–*19*). However, the precise mechanisms that regulate potentiation of specific synapses, depression of other synapses, and the co-operative reorganization of synaptic strength on individual neurons to mediate functional plasticity *in vivo*, remain unexplored.

In particular, it has been pointed out that the joint action of Hebbian and homeostatic plasticity in synaptic ensembles could impede the formation of new functions by cancelling each other (*19*). Alternatively synaptic potentiation at specific dendritic locations could be coordinated with heterosynaptic depression of nearby synapses within short stretches of the same dendrite in order to co-operatively implement functional plasticity of single cell responses. Such local plasticity could be particularly effective at reinforcing selective responses in dendritic compartments displaying functionally intermixed synaptic inputs. However, it is unknown whether locally coordinated synaptic plasticity occurs during physiological conditions *in vivo*, and whether it has a role in shaping neuronal responses. Here we have addressed these questions by using visual-optogenetic pairing to induce receptive field plasticity in single visual cortex neurons of awake mice, and asking how such plasticity is implemented by local Hebbian potentiation and heterosynaptic depression of identified synapses on neuronal dendrites.

## Induction of receptive field plasticity in single-neurons of awake mice

Classical *in vivo* plasticity induced by sensory deprivation or enrichment (*2*) inevitably results in large-scale polysynaptic functional and structural changes. To isolate synaptic changes in single neurons *in vivo*, we first developed a controlled paradigm for inducing plasticity at identified synapses on single neurons in the primary visual cortex (V1) of awake juvenile mice (P28-P35), using a pairing protocol where individual neurons were forced to fire action potentials soon after presentation of a target visual stimulus (*20*–*22*). A sparse noise stimulus ensemble (white squares on a 4 by 5 grid, see Fig. 1A and Materials and Methods) was used to measure single-cell receptive fields and to induce plasticity. Pairing visual stimuli presented at a target location with channelrhodopsin-2 (ChR2) driven spiking of a single neuron allowed us to increase the strength of a subset of excitatory synapses at spines on the neuron - as evident by a shift in the neuron’s receptive field towards the target location - and examine the effects on the function and structure of these and a wide range of other spines (Fig. 1B). First, we characterized the population excitatory synaptic inputs to V1 layer 2/3 neurons in response to sparse noise stimuli using whole-cell recordings in voltage-clamp mode (Fig. 1C). Excitatory subthreshold receptive fields were large, but with centroids that could be clearly identified (Fig. 1D-E). The EPSC evoked by the sparse noise stimuli typically displayed two components corresponding to stimulus onset and offset respectively (Fig. 1F). The response at stimulus onset was sharp, unimodal and lasted for about 200 ms with a peak response at 65-130 ms (mean 96.3±7.4 ms, n=8 neurons) after stimulus onset. Based on previous studies (*21*, *22*), we reasoned that pairing a visual stimulus at a target location with ChR2 activation of a postsynaptic neuron at a sufficient latency within the onset EPSC response (150 ms after stimulus onset) would be an effective pre-leading-post (synaptic input-action potential) protocol for inducing Hebbian potentiation of excitatory synapses responding to the target stimulus, and hence for inducing receptive field shifts at the soma.

**Fig. 1.**
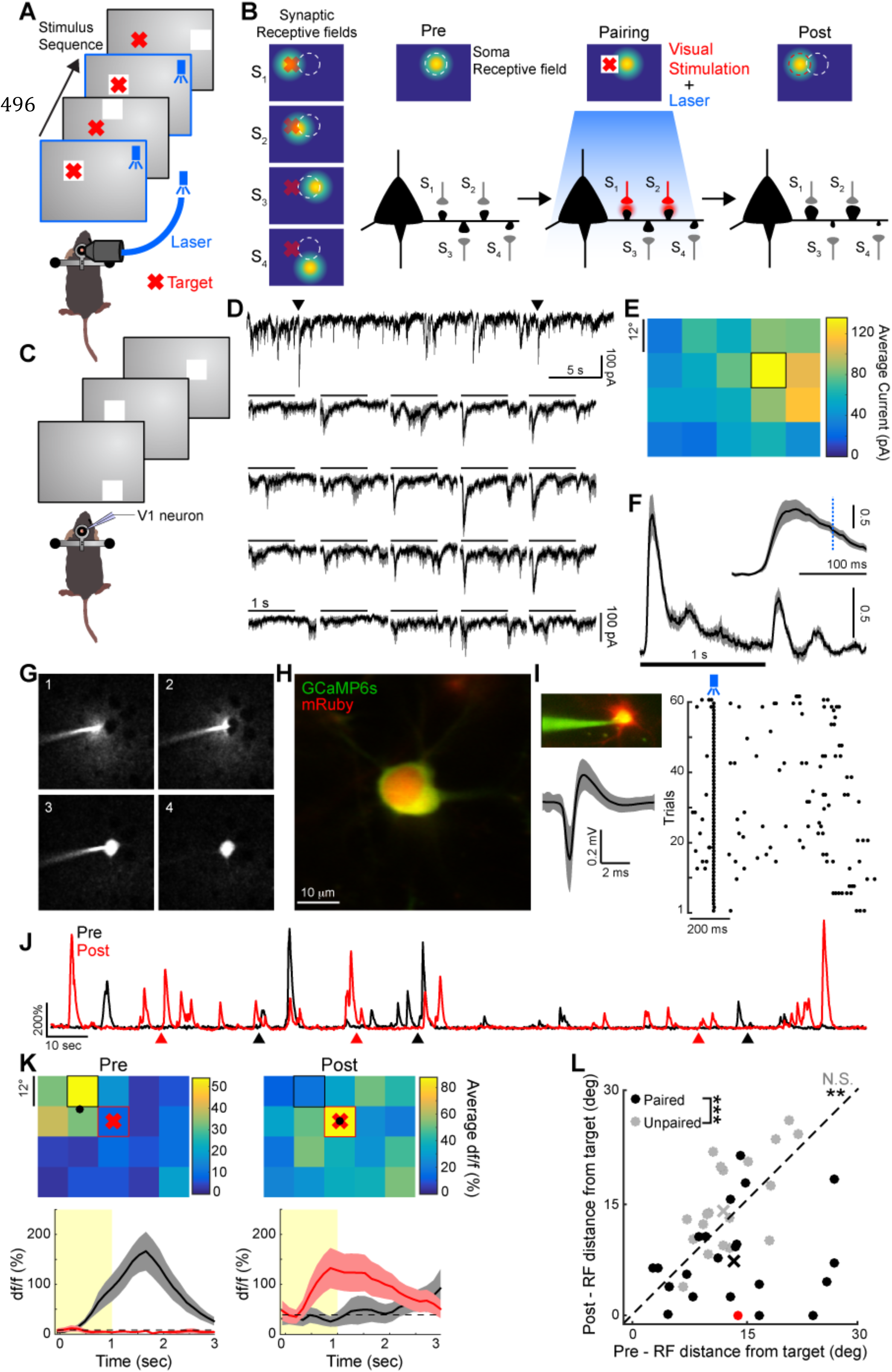
Functional plasticity induced in single V1 neurons of awake mice. (**A**) Schematic of the protocol used to induce plasticity. Whenever a stimulus (white square) appeared at a target location (red cross), ChR2 expressing neurons were stimulated with a blue laser pulse. (**B**) Illustration of the effect of the pairing protocol on a neuron’s receptive field and its dendritic spines. Left: Layer 2/3 excitatory neurons in V1 receive synaptic inputs (S_1_-S_4_) at spines from diverse local neurons (synaptic receptive fields). The soma receptive field, depicted above the neuron, represents the integration of these synaptic inputs. Here we illustrate the scenario where S_1_ and S_2_ receive inputs from cells with receptive fields close to the stimulated target location. Middle: Schematic of the plasticity protocol where visual target stimulation (red cross on white square) is repeatedly paired with ChR2 neuronal stimulation (blue). Right: After plasticity induction, the receptive field of the neuron shifts towards the target location. This is achieved through specific potentiation of spines with receptive fields close to the target and depression of other spines. The initial and final soma receptive field locations are shown as dashed white and red circles respectively. (**C**) Whole-cell recording of single neuron receptive field using sparse noise stimuli. (**D**) Top: example current trace of a neuron recorded in voltage-clamp mode at -70mV. Arrows indicate onset of preferred stimulus. Bottom: Average EPSC recorded for the same neuron for each stimulus location on a 4x5 grid. Scale bars indicate stimulus duration. Shaded gray areas: s.e.m. (**E**) Spatial ON receptive field obtained by averaging currents elicited by stimulus onset for each stimulus position (window: 50-150 ms). Black square indicates preferred stimulus position. (**F**) EPSC (converted to z-score) averaged over all neurons and all stimulus positions (n=8 neurons). Shaded area: s.e.m. Inset: expanded view of the first 200 ms. The dashed blue line indicates timing of ChR2 stimulation for pairing. (**G**) Snapshots showing single-cell approach (1-2) and subsequent electroporation *in vivo* (3). Cell body is left intact after withdrawal of the pipette (4). (**H**) Example of a neuron expressing GCaMP6s-P2A-mRuby and ChR2 48h after electroporation. (**I**) Loose-patch recording of an electroporated neuron expressing ChR2,showing responses to single blue light pulses. Left: Image of a patched neuron and averaged spike waveform (grey: standard deviation). Right: raster plot for several trials of blue light stimulation. (**J**) Example calcium df/f traces obtained from soma imaging before (black) and after (red) visual-ChR2 stimulation pairing. Arrows indicate visual stimuli, presented pre-pairing at the preferred location (black) and post-pairing at the target location (red). (**K**) Top: Receptive fields corresponding to the traces in **J**. Black and red squares indicate preferred stimulus location before and after pairing respectively. Black dots indicate center-of-mass and red crosses indicate target stimulus location for pairing. Bottom: Response time course for the preferred stimulus location before (black) and after pairing (red). Shaded area: s.e.m. Yellow areas indicate stimulus duration. (**L**) Comparison of the distance between target and receptive field center-of-mass before and after pairing (black, n=22 neurons with significant visual responses before and after pairing out of 94 electroporated cells, N=23 mice, paired Wilcoxon test, **p<0.01). The red dot indicates the example shown in panel **K** and the black cross indicates the average shift. Control experiments where ChR2 activation was not paired with the target stimulus are shown in gray (n=21 neurons out of 91, N=11 mice, paired Wilcoxon test, p=0.06, N.S. Not significant). Distribution of receptive field shifts were significantly different between these two populations (unpaired Kruskal-Wallis test, ***p<0.001)

To reliably induce and measure receptive field plasticity of single neurons over an extended period of time, we recorded visually-evoked Ca^2+^ transients of these neurons and optogenetically manipulated their firing. Using single-cell electroporation, we delivered two plasmids into single neurons encoding respectively the calcium indicator GCaMP6s (*23*) with mRuby fluorescent protein, and ChR2 (*24*) fused with a mCherry fluorescent protein (Fig. 1G,H, see Materials and Methods). Expression of ChR2 allowed precise and reliable control of neuron activity in awake mice with on average a single spike evoked for each blue light pulse (Fig. 1I, n=7 neurons, number of spikes/pulse = 1.2±0.32). Prior to inducing receptive field plasticity in single cells, we used the GCaMP6s signal to map the receptive field measured at the soma (Fig. 1J-K,Fig. S1A-B). Based on the spatial distribution of the receptive field we determined a target stimulus location in the vicinity of the peak response location. Repeated presentations of the target stimulus were then paired with single blue light pulses delivered to the cortex to elicit spikes in the imaged neurons (60 pairings, see Materials and Methods). A distractor stimulus was presented in-between each pairing to prevent fixation at the target location (Fig. 1A). The effect of this pairing was assessed by measuring the receptive field of the same neuron 1 to 2 hours after the end of the protocol. For most neurons, the receptive field center-of-mass shifted toward the target stimulus as a result of the pairing (Fig. 1K-L), indicating that the protocol successfully altered the neuron’s synaptic weights (Fig. 1B). These receptive field changes lasted for several hours and were not observed when ChR2 stimulation was not paired with the target stimulus (Fig. 1L). Moreover receptive field shifts could not be explained by changes in eye position statistics (Fig. S2) and did not induce noticeable functional changes in the surrounding network (Fig. S3). Successful receptive field shifts could also be achieved with long pairing times at low pairing rate (Fig. S4).

## Synaptic inputs are functionally heterogeneous along dendritic stretches

In order to understand how synapses on imaged neurons were affected by the plasticity protocol, we first defined responses of identified spines and their distribution on dendrites. GCaMP6s and mRuby signals were imaged in stretches of dendrites and spine-specific calcium responses were isolated (Fig. 2A-C). The shape and volume of spines were estimated based on the mRuby structural fluorescence signal. The fluorescent GCaMP6s activity in individual spines in response to sparse noise stimuli was used to measure input-specific receptive fields (see Materials and Methods and Fig. S1C-F). We found a heterogeneous distribution of spine receptive fields along the dendritic tree (Fig. 2D). This was assessed by comparing for each cell the distribution of receptive field center-of-mass distance between pairs of spines belonging to the same dendrite or to different dendrites (Fig. 2E). Although spines along the same dendritic stretches had significantly closer receptive fields than spines in different stretches, the two distributions overlapped considerably. Moreover, no clear spatial organization of receptive fields was found for spines lying along the same dendrite, indicating a functionally heterogeneous mixture of synapses, and of their presynaptic neurons, on dendrites (Fig. 2F-G).

**Fig. 2.**
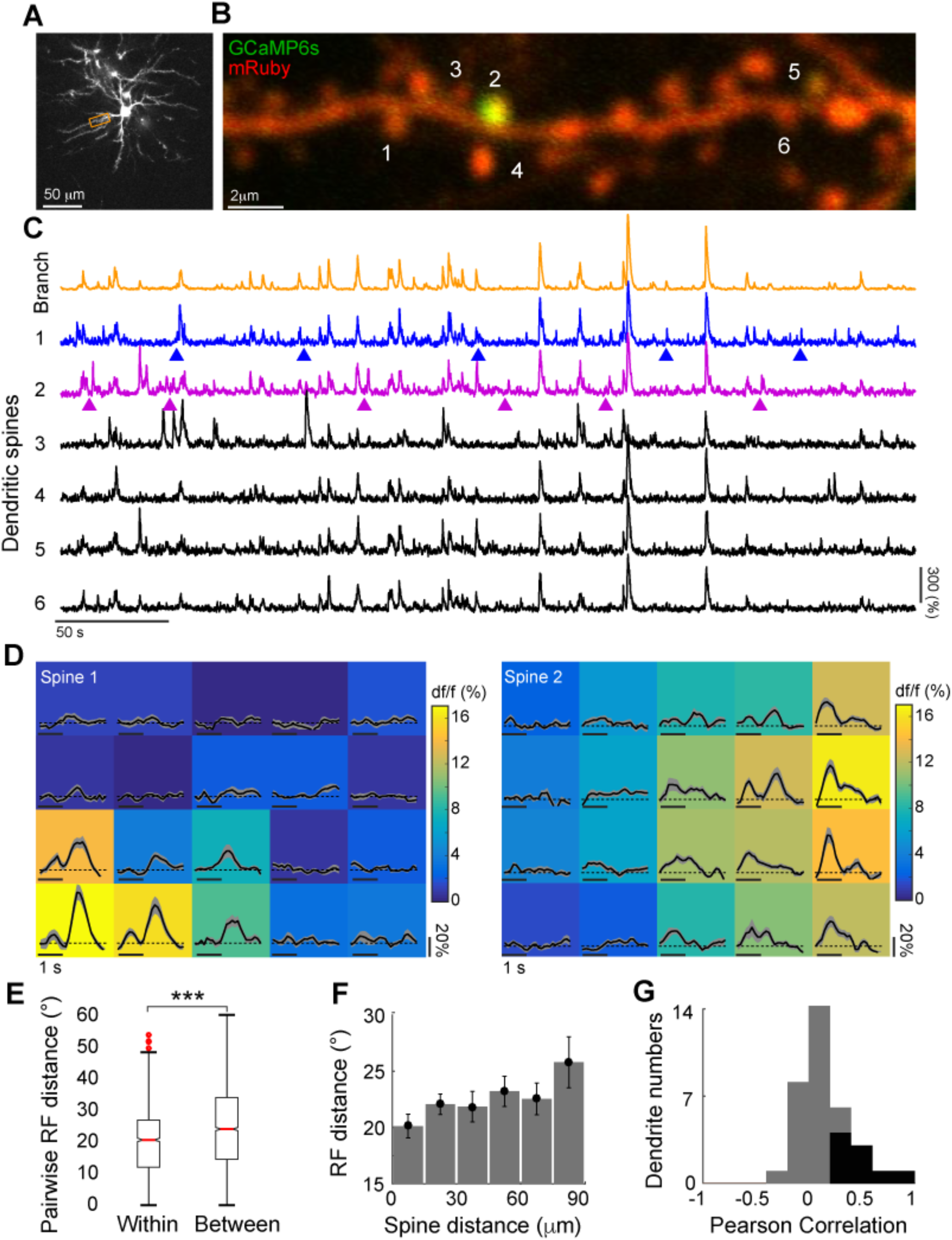
Synaptic inputs are functionally heterogeneous along dendritic stretches. (**A**) Z-stack projection of a neuron expressing GCaMP6s-P2A-mRuby. (**B**) Image of a dendritic stretch indicated by the orange rectangle in **A**. (**C**) Calcium fluorescent traces for representative spines indicated by numbers in **B**. Orange trace: signal imaged in the entire branch shaft. Arrows: onset of preferred stimuli for the first two spines (blue and purple). (**D**) Receptive field examples showing df/f responses for spines 1 and 2 in **B-C**. Average response traces are shown as black lines with s.e.m. in gray. Colormap indicates time-averaged magnitude for each location obtained over the whole response window. (**E**) Distribution of receptive field distances for pairs of spines within the same branch (n=1238 pairs) or between different branches (n=2498 pairs, unpaired Kruskal-Wallis test, ***p<0.001). (**F**) Relationship between spine distance on the dendrite and receptive field distance for pairs of spines in the same branch averaged over dendrites (n=34 dendrites, one-way ANOVA test, p=0.34). (**G**) Distribution of Pearson correlation coefficients between spine and receptive field distances measured in individual dendrites (n=34 dendrites, black indicates significant correlation with p<0.05).

## Hebbian potentiation and heterosynaptic depression in functionally heterogeneous dendritic spines

We next investigated the structural and functional basis at the synaptic level of receptive field plasticity at the soma (Fig. 1B). Spines receiving synaptic inputs during the narrow time window following target stimulus onset (Fig. 1F, inset) could be identified by averaging their calcium responses during the first second following stimulus onset (Fig. 2D). By repetitively pairing the activation of a subset of synapses to a somatic spike through ChR2 activation delivered 150 ms after stimulus onset, we specifically targeted these synapses for potentiation through spike-timing-dependent plasticity (*21*, *22*). To compare functional changes at the soma with plasticity in functionally identified spines, we used increased or decreased spine volume as a structural proxy for long-term potentiation (*25*–*27*) or depression (*27*) respectively. Because we used a short pairing protocol, we hypothesized that spines within the same dendritic branches that experience depression will do so through heterosynaptic plasticity acting on a slower time scale in response to local targeted potentiation. By comparing dendritic spines before and at several time points following the pairing protocol, we indeed observed bi-directional volume changes within the same dendritic stretch with seemingly different temporal dynamics (Fig. 3A). These changes were not the result of drift in the imaging planes (Fig. S5). Such structural long-term potentiation (sLTP) or depression (sLTD) in ChR2-expressing (ChR2+) neurons were compared to changes observed in control experiments where neurons did not express ChR2 (ChR2- neurons; Fig. 3B-C). We quantified changes in spine volume using the normalized difference between the integrated spine fluorescence signal relative to shaft before and after the pairing (see Materials and Methods). The normalized difference (δV) is contained in the range [-1,1] with positive values for spines that became larger after pairing and negative values for spines that became smaller (Fig. 3B). The proportion of spines for different volume change values revealed a balanced increase of sLTP and sLTD spines in ChR2+ neurons as measured at least 2 hours following the pairing (Fig. 3C). To assess the effect of the pairing protocol on spine structure above chance level, we defined a threshold for pairing-induced sLTP and sLTD in ChR2+ neurons based on the spine volume change distribution in ChR2- neurons (a δV value of ±0.25, corresponding to spines that exceeded the 97^th^ percentile of the ChR2- distribution; Fig. S6). By considering spines that became significantly potentiated or depressed more than two hours following pairing, we could backtrack their temporal evolution to the pairing protocol (Fig. 3D). We found that sLTP spines rapidly increased in volume right after pairing followed by a moderate increase over the following two hours. In contrast, sLTD spines showed a small decrease in volume soon after pairing that strongly declined over the course of the following two hours until the average volume change for both sLTD and sLTP spines became approximately balanced. These different temporal dynamics indicate that fast Hebbian potentiation is first induced in targeted spines closely followed by heterosynaptic depression of other spines acting on the timescale of minutes to hours. We then investigated coordination between sLTP and sLTD spines within individual dendrites by comparing their density and distribution (Fig. 3E). The density of sLTD spines was significantly correlated with, and was greater than, the density of sLTP spines (Fig. 3F). Because spine density is quite variable along dendrites, we examined the distribution of depressed spines around potentiated spines by computing the deviation of sLTD spine density from the mean as a function of distance from sLTP spines for each dendrite (see Materials and Methods). We reasoned that if sLTD spines were more likely to be found around sLTP spines, we would see a significantly larger density at short sLTP-sLTD distances. The pairing protocol indeed resulted in significantly larger sLTD spine density in the vicinity of sLTP spines (Fig. 3G).

**Fig. 3.**
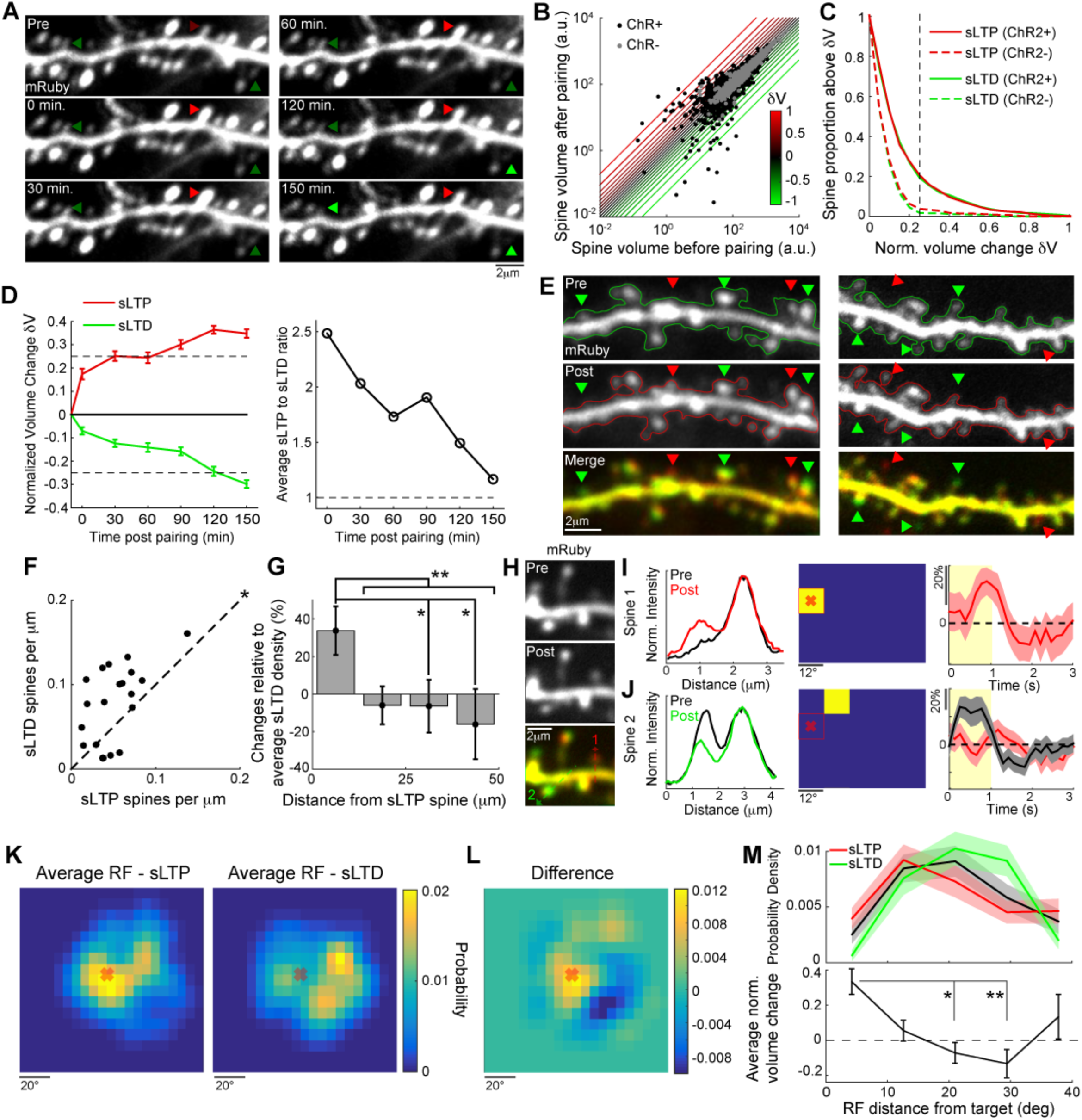
Hebbian potentiation and heterosynaptic depression in functionally
heterogeneous dendritic spines. (**A**) Time-lapse imaging of a dendritic stretch before and after pairing. Red and green arrows indicate example spines that undergo structural long-term potentiation (sLTP) and depression (sLTD) respectively. (**B**) Comparison of spine volumes before and more than 2 hours after pairing for ChR2+ (black, n=1987 spines) and ChR2-(gray, n=845 spines) neurons including newly formed or eliminated spines. The colored lines indicate domains of constant normalized volume change in the [-1,1] interval. (**C**) Proportion of sLTP (red) and sLTD (green) spines for different values of normalized volume change. Curves for ChR2- neurons are shown with dashed lines (variance F-test between volume change distributions for ChR2+ and ChR2- spine populations, p<0.001). Black dashed line indicates sLTP and sLTD plasticity threshold of δV>0.25 based on the ChR2- control data (Fig. S6). (**D**) Left: For all spines that were significantly potentiated (red, n=110) or depressed (green, n=98) at least 2 hours following the pairing, the average normalized volume change is shown for the preceding time points. Error bars: s.e.m. Dashed lines indicate threshold for sLTP and sLTD. Right: sLTP to sLTD volume change ratio for all time points following pairing. Dashed line indicates identical average volume change. (**E**) Examples of dendritic stretches where sLTP (red arrows) and sLTD (green arrows) spines are intermixed. Dendrite contours are depicted before and after pairing. (**F**) Comparison of sLTP and sLTD spine density per dendrite (n=20 dendrites; Pearson coefficient: 0.55, p<0.05; paired Wilcoxon test, * p<0.05). (**G**) For dendrites in **F**, sLTD spine density variation relative to the mean as a function of distance from sLTP spines in individual dendrites (average over n=103 sLTP spines with neighbor sLTD spines, one-way ANOVA test, p<0.01, unpaired Kruskal-Wallis test, **p<0.01 and *p<0.05 with Bonferroni correction). The distribution of sLTD spines in the immediate vicinity of sLTP spines showed a significantly larger density increase compared to longer distances. (**H**) Example of a dendritic segment where an enlarged spine is in close proximity to a depressed spine following the pairing. The bottom panel compares the dendrite before (green) and after (red) plasticity induction. (**I**) Left: Profiles indicated by the dashed red line in **H** (bottom) comparing spine size before (black) and after (red) pairing. Middle: receptive field of the corresponding spine (yellow square indicates location with significant responses and red cross indicates target position). Right: Response time course for the preferred stimulus. Yellow area indicates stimulus duration and shaded red areas s.e.m. (**J**) Same as **I** for the depressed spine in **H** indicated with a green dashed arrow. Right: Response time course for the preferred stimulus (gray) and for the target stimulus location (red). (**K**) Average normalized receptive field centered on target for spines that experienced sLTP (left, n=94 spines) and sLTD (right, n=87 spines). (**L**) Difference between the distributions in **K**. (**M**) Top: Distribution of receptive field distances from target for sLTP spines (red) and sLTD spines (green). The black distribution is obtained by randomly shuffling spine identity. Shaded areas indicate standard deviation. Bottom: Average normalized volume change as a function of receptive field distance from target for all spines that experienced plasticity (n=181 spines, one-way ANOVA test, p<0.01, * p<0.05 and ** p<0.01 with Bonferroni correction).

We investigated the functional signature associated with these structural changes to evaluate if they were consistent with receptive field plasticity measured at the soma. In particular, we hypothesized that sLTP spines should have their receptive field centers close to or overlapping the visual target as a consequence of Hebbian plasticity, whereas sLTD spines would have receptive field centers located away from the target as a consequence of heterosynaptic, potentially co-operative, plasticity. Spines with receptive fields overlapping the target stimulus indeed increased in volume (Fig. 3H,I), whereas spines with receptive fields centered away from the target were reduced in volume (Fig. 3H,J). These structural changes were accompanied by consistent changes in GCaMP6s signal amplitude (Fig. S7). To examine at the population level whether functional plasticity of neuronal receptive fields was accompanied by structural and functional changes in spines, we examined the receptive fields of all spines that underwent sLTP or sLTD, relative to the target stimulus location used for each neuron (Fig. 3K). The average receptive field for sLTP spines was sharp and centered on the target whereas the average receptive field for sLTD spines was distributed broadly away from and around the target. The difference between these average receptive fields (Fig. 3L) highlighted the observation that sLTP spines were specifically targeted by the pairing protocol whereas sLTD spines had receptive fields away from the target and resulted in response reduction around the target stimulus. Quantifying the spine volume difference for sLTP and sLTD spines as a function of their receptive field center distance from the target stimulus showed that as the distance from target increased, the effect on spine size shifted from sLTP to sLTD (Fig. 3M). The organized distribution of sLTP and sLTD spines, and the location of their receptive fields, demonstrates that targeted Hebbian potentiation accompanied by local heterosynaptic depression remodels synaptic inputs within functionally heterogeneous dendrites, to enable cell-wide functional receptive field plasticity.

## Arc and AMPAR dynamics in dendrites following receptive field plasticity

To assess mechanisms of functional and structural changes resulting from the pairing protocol, we electroporated AMPA receptor (AMPAR) subunit 1 tagged with a pH-sensitive form of GFP (Super Ecliptic pHluorin) SEP-GluA1 (*18*, *28*) into single neurons to restrict the signal to membrane-inserted receptors, together with ChR2 and a volume-filling marker DsRed2, and imaged neurons 5-10 days after electroporation. Because SEP-GluA1 utilizes GFP as indicator, and to determine a target stimulus for plasticity induction, we electroporated GCaMP6s into neighboring neurons (Fig. 4A) that shared a substantial proportion of their subthreshold receptive field (Fig. S8) – the target was placed near their receptive field. Several days after electroporation, a strong and distinct SEP fluorescence signal could be observed at the surface of individual spines (Fig. 4B, left) indicating the density of AMPARs inserted in the membrane. Comparing the normalized change in SEP-GluA1 density and volume in individual spines two or more hours following the short or long pairing protocol (Fig. S4), we found a significant positive correlation (Fig. 4C-E), indicating that positive or negative changes in volume reflected corresponding modifications of spine synaptic weight through AMPAR expression changes.

**Fig. 4.**
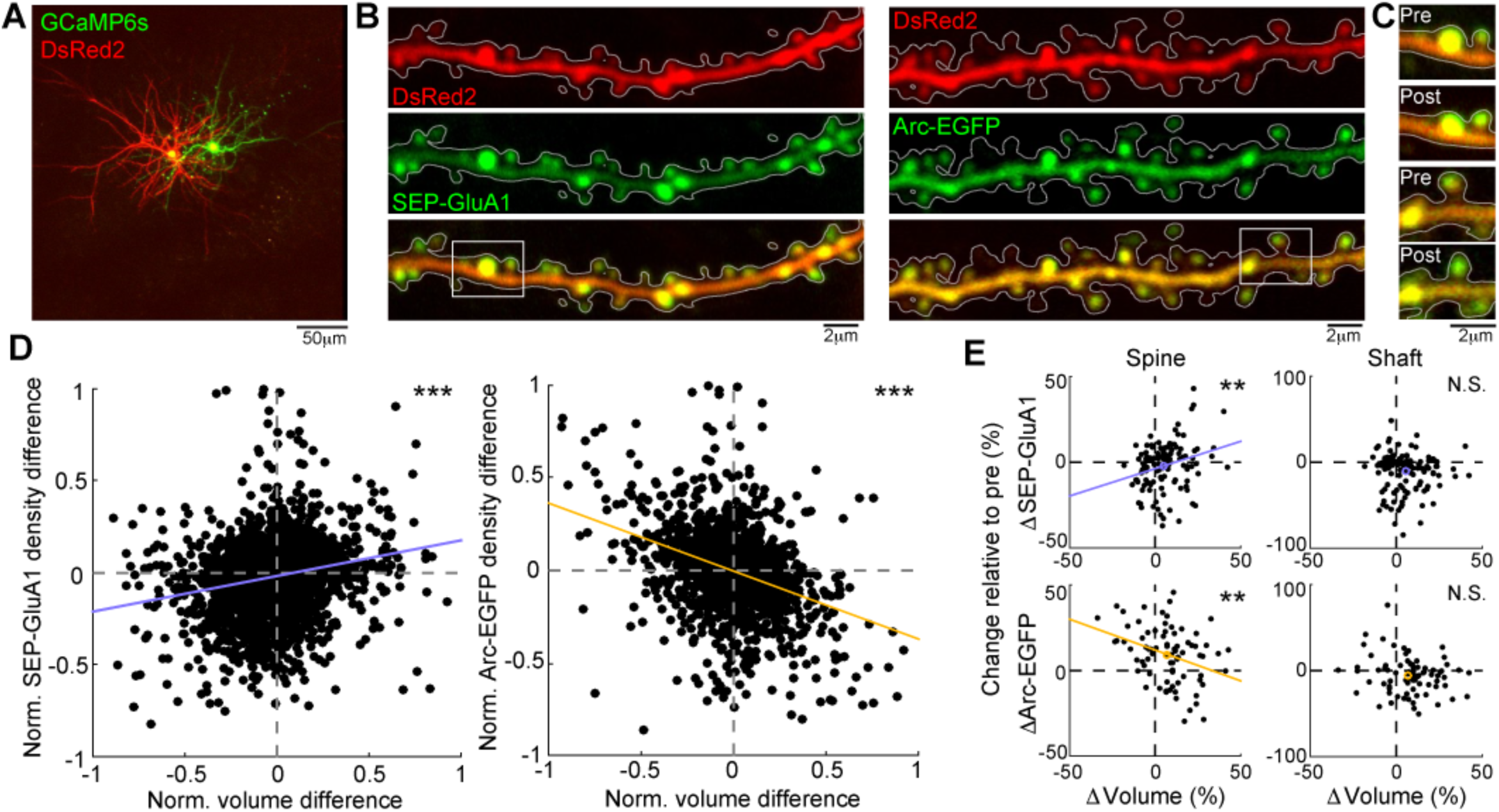
Arc and AMPAR trafficking in dendrites following receptive field plasticity. (**A**)Example V1 neuron expressing DsRed2 and SEP-GluA1 in close proximity to a neuron expressing GCaMP6s. (**B**) Example of dendrites expressing DsRed2 and SEP-GluA1 (left) or Arc-EGFP (right). Dendrite contours are shown in white. (**C**) Example of spine changes before and after pairing for SEP-GluA1 (top) or Arc-EGFP (bottom) corresponding to the white rectangles in **B**. (**D**) Comparison of the normalized change in volume and SEP-GluA1 (left) or Arc-EGFP (right) density for individual spines (SEP-GluA1: n=4354 spines from 17 neurons, N=12 mice, Pearson coefficient=0.22, *** p<0.001; Arc-EGFP: n=1719 spines from 16 neurons, N=8 mice, Pearson coefficient=-0.37, *** p<0.001). These panels combine data obtained with the short pairing protocol (SEP-GluA1: n=2046 spines, Pearson coefficient=0.28, p<0.001; Arc-EGFP: n=262 spines, Pearson coefficient=-0.32, p<0.001) and the long pairing protocol (Fig. S4, SEP-GluA1: n=2308 spines, Pearson coefficient=0.16, p<0.001; Arc-EGFP: n=1457 spines, Pearson coefficient=-0.4, p<0.001). The purple and orange lines indicate linear fits for SEP-GluA1 and Arc-EGFP respectively. (**E**) Average changes in SEP-GluA1 (top, n=122 dendrites) and Arc-EGFP (bottom, n=81 dendrites) density relative to initial density as a function of average spine volume change relative to initial volume for individual dendrites. The left and right plots show the comparison for SEP-GluA1 or Arc-EGFP signal average over all spines and in the dendritic shaft respectively. Only signals in spines correlate with structural plasticity (Pearson coefficient: 0.27, ** p<0.01 for SEP-GluA1 and -0.29, ** p<0.01 for Arc-EGFP, N.S. Not Significant). The purple and orange lines indicate best linear fits and empty circles indicate average values.

The immediate early gene Arc has been previously shown to be involved in AMPAR endocytosis (*29*) and in mediating homeostatic plasticity in cell cultures (*30*). Recent work has suggested that Arc preferentially interacts with the β isoform of CaMKII and acts as an inverse tag of plasticity (*31*) that could potentially act in local dendritic segments to mediate heterosynaptic depression (*32*). However, the single-spine dynamics of Arc *in vivo* in response to plasticity induction remain unexplored. We employed a monomeric EGFP-tagged Arc (mEGFP-Arc) probe under the control of the Arc promoter to study the molecular dynamics of Arc following the pairing protocol (*31*) (Fig. 4B, right). The accumulation of Arc in dendritic spines becomes strongly significant about 2 hours following plasticity induction *in vitro* (*31*). Interestingly, we found that Arc-EGFP changes during volume changes were complementary to and mirrored changes in AMPAR density (Fig. 4C-E). Thus, Arc expression density was increased in sLTD spines and decreased in sLTP spines, supporting the idea that Arc acts as an inverse tag on spines that did not previously experience potentiation and therefore could be subject to heterosynaptic depression. We further tested if the level of Arc-EGFP and SEP-GluA1 modulation was dendrite-specific. We found that the average spine volume change for individual dendrites was correlated with the average change of spine SEP-GluA1 and anti-correlated with the average change of spine Arc-EGFP (Fig. 4E). These correlations were not found for Arc-EGFP and SEP-GluA1 signals in the dendritic shaft.

## Role of Arc in regulating AMPAR distribution and receptive field plasticity

To further examine the role of Arc in mediating spine-specific heterosynaptic depression *in vivo*, we delivered small hairpin RNA (shRNA, see Materials and Methods and Fig. S9 and S10) to deplete Arc in single neurons. This plasmid was electroporated together with SEP-GluA1 to track surface AMPAR distribution (Fig. 5A). We imaged neurons 5-10 days after electroporation to study the long-term effects of Arc depletion. Neurons in which Arc was knocked down displayed dendrites populated with mature spines filled with SEP-GluA1 compared to control neurons with unaltered Arc expression (Fig. 5B). The distribution was skewed with a large variance in the control condition, indicating a more heterogeneous distribution of surface AMPAR density as compared to the knock down condition, while the latter had a lower ‘hot spot index’ (see Materials and Methods) indicating more closely located spines with high surface SEP-GluA1 density (Fig. 5B, inset). The functional consequence of abnormal AMPAR distribution was assessed by using the pairing protocol in Arc knock down neurons expressing GCaMP6s and ChR2 (Fig. 5C). We quantified the change in calcium activity following pairing by measuring average amplitude and frequency of GCaMP6s transients for neurons with unaltered Arc or Arc knock down (Fig. 5D-E). Neurons with normal Arc expression showed no changes in the average amplitude of GCaMP6s transients but displayed a significant reduction of transient frequency. We posit that this is caused by a strong synaptic depression in response to the pairing protocol to balance the newly potentiated spines. In contrast, neurons with reduced Arc expression displayed increased transient frequency after the pairing protocol (Fig. 5E). These data suggest that co-operation between synaptic potentiation and depression is altered in neurons lacking Arc expression, likely causing reduced AMPAR endocytosis. We then compared the distance between neuronal receptive field centers and target before and after the pairing, and found that knock down of Arc prevented displacement of receptive fields toward the target (Fig. 5F), consistent with impaired functional plasticity (*33*). Functional and structural imaging of spines indicated that the lack of neuronal receptive field plasticity could potentially be explained by reduced functional segregation between potentiated and depressed spines (Figs. S10 and S11). [Note that we did not find any changes in spine density following the pairing protocol in any of these conditions (Fig. S12)]. Thus, Arc is critical for altering AMPAR expression at synapses and driving a mechanism that underlies receptive field plasticity of neurons.

**Fig. 5.**
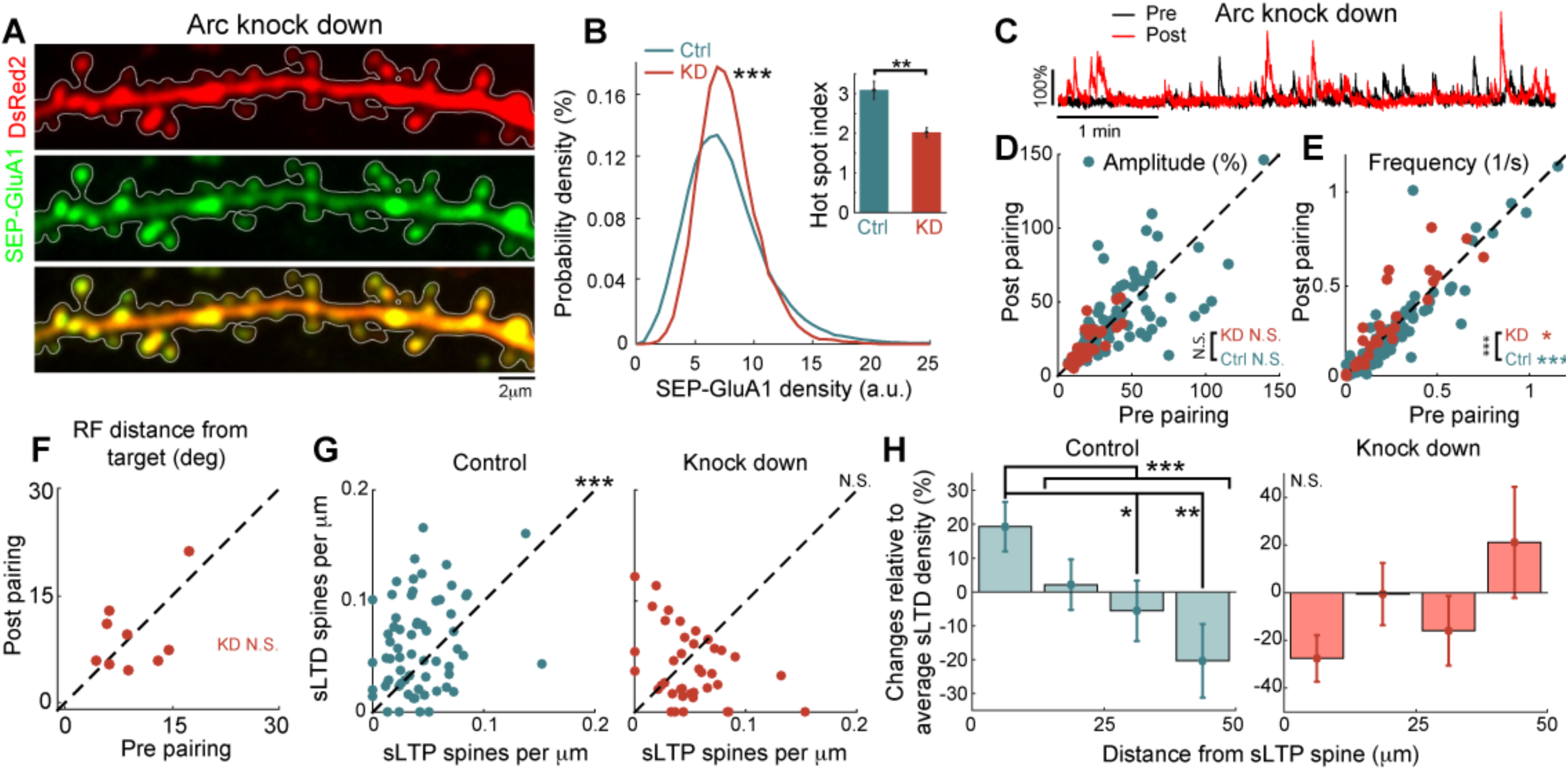
Role of Arc in regulating AMPAR distribution at spines and receptive field plasticity of single V1 neurons. (**A**) Example of a dendrite expressing SEP-GluA1 and ArcshRNA (Arc knock down) fused with DsRed2. (**B**) Distributions of SEP-GluA1 density for individual spines where Arc is endogenously expressed or knocked down (n=992 spines from 3 neurons for KD, n=5302 spines from 17 neurons for Ctrl, Kruskal-Wallis test, *** p <0.001). Inset: hot spot index measured for individual dendrites in these conditions (n=18 dendrites for KD and n=123 dendrites for control, unpaired Kruskal-Wallis test, **p<0.01). (**C**) Example of calcium traces before and after pairing for a neuron with Arc KD. (**D**) Comparison for individual cells of the average GCaMP6s transient amplitude in response to sparse noise before and after pairing for control (turquoise, n=94 neurons, N=23 mice) or Arc KD (brown, n=27 neurons, N=7 mice) conditions (paired Wilcoxon test, N.S. Not significant, comparison between KD and Ctrl conditions was performed with an unpaired Kruskal-Wallis test between distributions for post-pre differences). (**E**) Same as in **D** but comparing transient frequency (paired Wilcoxon test, p<0.001 for control and p<0.05 for KD; unpaired Kruskal-Wallis test for comparing both populations, p<0.001). (**F**) Comparison of the distance between target and receptive field center-of-mass before and after pairing for the KD condition (n=9 neurons with significant visual responses before and after pairing out of 27 electroporated cells, N=7 mice, paired Wilcoxon test, N.S. not significant). (**G**) Comparison of sLTP and sLTD spine density per dendrite for control (left, n=66 dendrites; Pearson coefficient: 0.31, p<0.05; paired Wilcoxon test, *** p<0.001) and Arc KD neurons (right, n=39 dendrites; Pearson coefficient: -0.37, p<0.05; paired Wilcoxon test, N.S. Not Significant) conditions. (**H**) For dendrites in **G**, sLTD spine density variation relative to the mean as a function of distance from sLTP spines in individual dendrites for the control (left, average over n=253 sLTP spines with neighbor sLTD spines, one-way ANOVA test, p<0.001, unpaired Kruskal-Wallis test, **p<0.01 and *p<0.05 with Bonferroni correction) and Arc KD conditions (right, average over n=142 sLTP spines with neighbor sLTD spines, one-way ANOVA test, p=0.3). Dendrites in the control condition either express a scrambled Arc shRNA plasmid fused with DsRed (n=46 dendrites) or GCaMP6s-P2A-mRuby (n=20 dendrites).

We then investigated whether coordinated spine potentiation and depression within individual dendrites was altered in the absence of Arc. In contrast to neurons with normal Arc expression, the density of sLTP spines was not significantly different from the density of sLTD spines for neurons with Arc knock down (Fig. 5G). Furthermore, in dendrites with normal Arc expression, the pairing protocol resulted in significantly larger sLTD spine density in the vicinity of sLTP spines (Fig. 5H, left) whereas this effect was absent when Arc was knocked down, (Fig. 5H, right) indicating impaired spatial organization of heterosynaptic plasticity around potentiated spines. In all these conditions we did not observe any specific organization of potentiated spines along the dendrite (Fig. S13). Thus, Arc helps organize the distribution of potentiated and depressed spines that underlies neuronal plasticity.

## Spine-specific Hebbian potentiation and heterosynaptic depression at eye opening following monocular deprivation

Finally, we examined whether Arc-mediated spine-specific heterosynaptic interactions, specifically the paired expression of sLTP and sLTD in nearby spines, occurs under physiological conditions where neurons are not manipulated with optogenetics. V1 neurons show reduced responses from a deprived eye following monocular deprivation, followed by recovery of responses when the deprived eye is re-opened (Fig. 6A); loss of Arc abolishes the effects of monocular deprivation or visual experience (*33*). We imaged V1 neurons in the binocular zone following 4-5 days of monocular deprivation before and after eye opening during the critical period for ocular dominance plasticity (*2*) (Fig. 6A-C), to assess how an increase in synaptic drive from the reopened eye remodeled spines. The ocular dominance index (ODI) was measured in neurons expressing GCaMP6s-P2A-mRuby (Fig. 6D), at the soma and at spines, by presenting drifting gratings to each eye separately (see Materials and Methods). Layer 2/3 neurons responded to visual stimuli in both eyes (Fig. 6E) and had dendrites where spines with preference for one eye or the other were intermingled (Fig. 6F-G). Following monocular deprivation, the population distribution of ocular dominance index for all imaged spines with visual responses was biased toward the contralateral eye, though a substantial proportion of spines had significant ipsilateral responses. We then assessed the level of structural plasticity several hours following eye opening, hypothesizing that specific potentiation of ipsilateral eye dominated spines would be accompanied by local heterosynaptic depression of contralateral eye dominated spines (Fig. 6I). We found a significant increase of potentiated and depressed spines as compared to age-matched control neurons where no monocular deprivation was performed (Fig. 6J). For neurons expressing DsRed2 and Arc-EGFP we found a negative correlation of Arc-EGFP density changes with spine volume changes, comparable to that obtained with the pairing protocol (Fig. 6K; cf. Fig 4D). Reopening the deprived eye led not only to potentiated spines but also to a significantly larger density of depressed spines in individual dendrites (Fig. 6J,L). Similar to the pairing protocol, sLTD spines were found in significantly higher density around sLTP spines (Fig. 6M). Finally, we related spine volume changes to their functional properties. Spines with ODI biased toward the ipsilateral eye (ODI<0) experienced strong potentiation whereas spines with responses biased towards the contralateral eye were more subject to depression (Fig. 6N-O). Thus, spine-specific plasticity accompanies recovery of V1 neurons from monocular deprivation, and likely involves coordinated spatial interactions between spines expressing Hebbian potentiation and heterosynaptic depression.

**Fig. 6.**
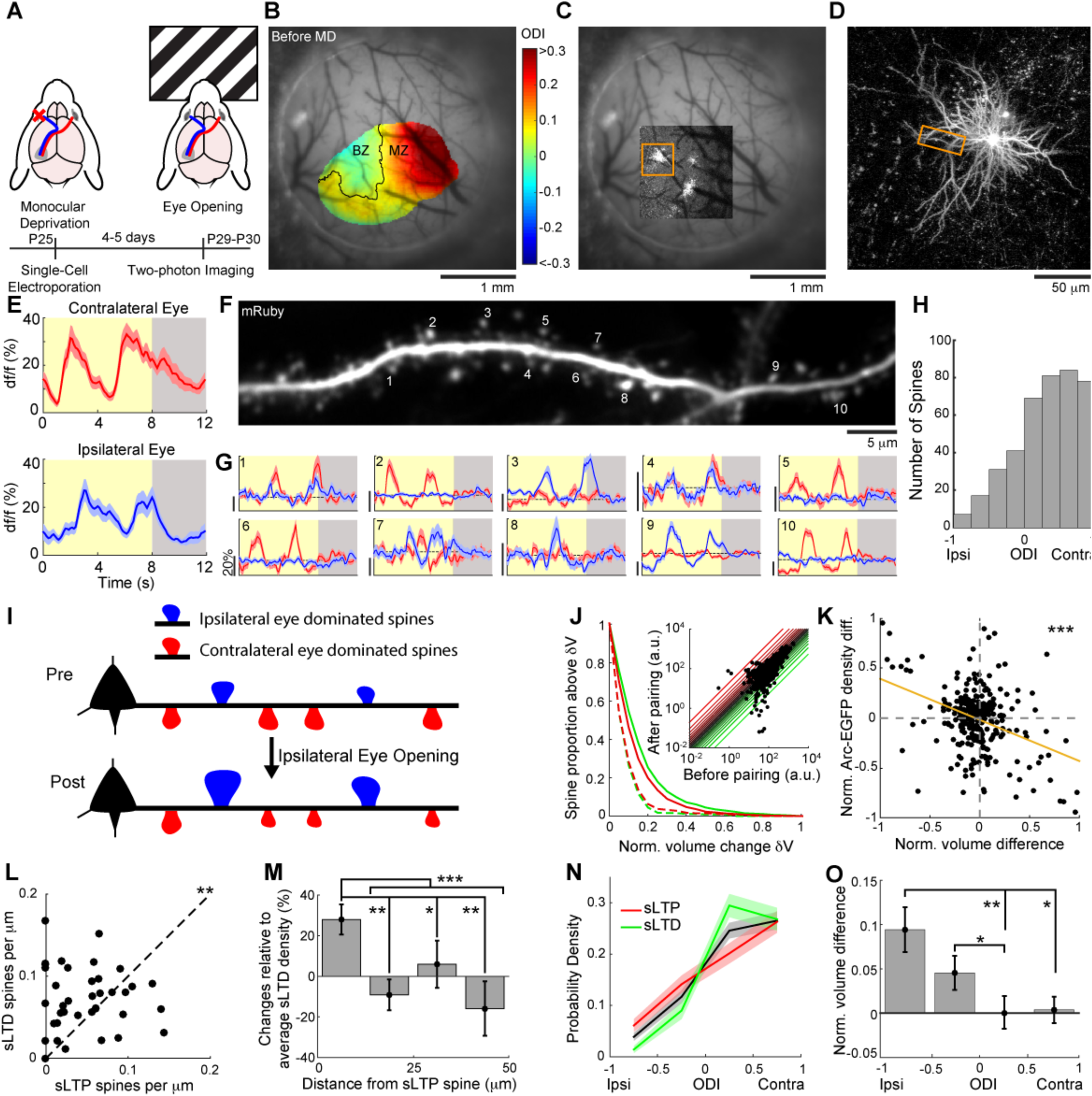
Spine-specific Hebbian potentiation and heterosynaptic depression at eye opening following monocular deprivation. (**A**) Schematic of the experiment. The ipsilateraleye (blue) was suture for a period of 4-5 days, starting at P25, during which the contralateral eye (red) was left open. On the last day, structural and functional imaging was performed before and after eye opening. (**B**) Ocular dominance domains obtained with intrinsic signal optical imaging. The color code indicates ODI and the border between the monocular zone (MZ) and the binocular zone (BZ) is indicated. (**C**) Low magnification two-photon image superimposed on the cranial window image. (**D**) Z-stack maximum projection of the neuron indicated by an orange rectangle in **C**. (**E**) Average df/f somatic traces of the cell in **D** in response to drifting gratings for the contralateral (top, red) and ipsilateral (bottom, blue) eyes. Shaded blue and red areas: s.e.m. Yellow areas indicate stimulus duration and gray areas blank screen presentation. (**F**) Magnification of the dendritic segment indicated by an orange rectangle in **D**. (**G**), Response df/f traces for individual spines indicated with numbers in **F**. Vertical scale bars: 20%. (**H**) Distribution of ODI values for all visually-responsive spines (n=408 spines from 14 neurons, N=3 mice). (**I**) Schematic of the spine size measurements and hypothesis. At eye opening spines receiving ipsilateral eye dominated synaptic inputs are expected to enlarge through Hebbian plasticity. As a consequence, nearby spines that receive contralateral eye dominated inputs are hypothesized to shrink through local heterosynaptic interaction. (**J**) Proportion of spines for different volume change thresholds (sLTP in red and sLTD in green, n=3028 spines from 13 neurons, N=5 mice). Dashed curves: baseline structural changes as in Fig. 3C (unpaired variance F-test with control condition, p<0.001). Inset: Comparison of spine volumes measured before and after eye opening. (**K**) Comparison of the normalized change in volume and Arc-EGFP density for individual spines (n=288 from 7 neurons, N=3 mice, Pearson coefficient=-0.38, *** p<0.001). The orange line indicates linear fit. (**L**) Comparison of sLTP and sLTD spines density for individual dendrites (n=39 dendrites, paired Wilcoxon test, ** p<0.01). (**M**) For dendrites in **L**, sLTD spinedensity variation relative to the mean as a function of distance from sLTP spines in individual dendrites (average over n=145 sLTP spines with neighbor sLTD spines, one-way ANOVA test, p<0.001, unpaired Kruskal-Wallis test, **p<0.01 and *p<0.05 with Bonferroni correction). (**N**) ODI distributions for sLTP spines (red) and sLTD spines (green). Black distributions are obtained by randomly shuffling spine identity. Shaded areas indicate standard deviation. (**O**) Normalized volume difference as a function of ODI for all spines (n=408 spines from 14 neurons, N=3 mice, one-way ANOVA test, p<0.01, * p<0.05 and ** p<0.01 with Bonferroni correction). Error bars: s.e.m.

## Discussion

We demonstrate here that coordinated Hebbian potentiation and heterosynaptic depression within local stretches of dendrites is a key mechanism underlying functional plasticity of single V1 neurons in awake animals *in vivo*. Potentiation of synapses with receptive fields overlying a target visual location, or driven by a newly opened eye after deprivation, is locally coordinated with depression of adjacent synapses with receptive fields that are off-target or driven by the other eye. Together, these synaptic changes co-operatively drive functional plasticity of single neuron responses and shift their receptive field towards the target visual location, or their ocular dominance towards the opened eye. These results provide a new understanding of the mechanisms underlying the functional and structural reorganization of dendritic spines following learning (*3*–*5*) and sensory loss or deprivation (*34*–*36*). Furthermore, we show that Arc-mediated heterosynaptic plasticity acting at the dendritic level can organize functional plasticity in neurons that have heterogeneous functional synaptic inputs. This mechanism can potentially confer on individual neurons the capacity to co-operatively alter functional properties that are locally distributed in different domains of their dendritic tree (*37*) (Fig. S14).

Locally coordinated potentiation of individual synapses together with depression of adjacent synapses contrasts sharply with cell-wide global homeostatic plasticity (*6*), which was proposed primarily as a mechanism acting over long timescales (hours to days) by which neuronal firing rates are maintained within a narrow physiological range. The role of local heterosynaptic depression overlaid on heterogeneous synaptic inputs, acting together with sparsely distributed potentiated spines (*38*), has been proposed through computational models (*19*) to produce rich synaptic integration at the single-cell level. Our results are in line with these predictions and show the functional relevance of this inverse synaptic tagging mechanism in regulating functional neuronal plasticity within hours in the intact brain. The spatio-temporal profile of the structural changes associated with heterosynaptic depression observed in our data (~10μm, ~2hours) is comparable to a previous study performed in hippocampal slices (*13*), possibly indicating a common mechanism (*32*).

Our findings potentially provide a functional explanation for the enigmatic targeting dynamics of Arc (*39*). Early studies of Arc found that domain-specific NMDA-dependent potentiation of synaptic inputs leads to augmented localization of Arc mRNA and proteins near the site of potentiation (*40*). Our findings, in conjunction with the role of Arc in mediating AMPAR endocytosis (*29*) in inactive spines (*31*), clarify its role in the spatial orchestration of LTD and LTP in functionally heterogeneous dendrites *in vivo*, whereby Arc is targeted near potentiated spines to weaken neighboring synaptic inputs through local heterosynaptic interactions.

## Acknowledgements

We thank Rachael Neve, Benjamin Bartelle and Alan Jasanoff for their assistance with plasmid preparation and testing. We thank Jeremy Petravicz for performing the eyelid sutures for MD experiments, Murat Yildirim for providing two-photon point spread function measurements and Keji Li for his assistance with optical intrinsic imaging. We thank Olivier Marre, Johannes Mayrhofer and Celia Gasselin for their comments on the manuscript. We thank Vivek Jayaraman, Rex A. Kerr, Douglas S. Kim, Loren L. Looger and Karel Svoboda from the GENIE Project, Janelia Farm Research Campus, Howard Hughes Medical Institute (HHMI) for the distribution of GCaMP6. This work was supported by Marie Curie postdoctoral fellowship FP7-PEOPLE-2010-IOF (S.E.B.), Human Frontier Science Program Long-Term Fellowship (J.P.K.I), AMED-CREST (H.B.) and KAKENHI grants (H.O., H.B.), and NIH grants NS090473 and EY007023; NSF grant EF1451125; the Simons Center for the Social Brain; and the Picower Institute Innovation Fund (M.S.).

## Author Contribution

S.E.B., J.P.K.I. and M.S conceived experiments. S.E.B. performed single-cell electroporation. S.E.B. and J.P.K.I. performed surgeries and carried out *in vivo* experiments. S.E.B performed data analysis. M.S. and J.P.K.I. contributed to analysis of experiments. V.B-P. performed intracellular recordings *in vivo* and eye tracking controls. H.O. and H.B. designed and provided the Arc-EGFP plasmid. S.E.B., J.P.K.I., V.B-P. and M.S. wrote the paper.

## Supplementary Materials

Materials and Methods

Figure S1-S14

